# Unique vulnerability of *RAC1*-mutant melanoma to combined inhibition of CDK9 and immune checkpoints

**DOI:** 10.1101/2023.06.27.546707

**Authors:** Alexa C. Cannon, Konstantin Budagyan, Cristina Uribe-Alvarez, Alison M. Kurimchak, Daniela Araiza-Olivera, Kathy Q. Cai, Suraj Peri, Yan Zhou, James S. Duncan, Jonathan Chernoff

**Affiliations:** Cancer Signaling and Microenvironment Program, Fox Chase Cancer Center, Philadelphia, PA; Drexel University College of Medicine, Philadelphia, PA; Histopathology Facility, Fox Chase Cancer Center, Philadelphia, PA; Biostatistics-Bioinformatics, Fox Chase Cancer Center, Philadelphia, PA; Current Affiliation: Merck, Bioinformatics Oncology Discovery, Boston, MA

## Abstract

*RAC1^P29S^* is the third most prevalent hotspot mutation in sun-exposed melanoma. *RAC1* alterations in cancer are correlated with poor prognosis, resistance to standard chemotherapy, and insensitivity to targeted inhibitors. Although *RAC1^P29S^* mutations in melanoma and *RAC1* alterations in several other cancers are increasingly evident, the RAC1-driven biological mechanisms contributing to tumorigenesis remain unclear. Lack of rigorous signaling analysis has prevented identification of alternative therapeutic targets for *RAC1^P29S^*-harboring melanomas. To investigate the RAC1^P29S^-driven effect on downstream molecular signaling pathways, we generated an inducible RAC1^P29S^ expression melanocytic cell line and performed RNA-sequencing (RNA-seq) coupled with multiplexed kinase inhibitor beads and mass spectrometry (MIBs/MS) to establish enriched pathways from the genomic to proteomic level. Our proteogenomic analysis identified CDK9 as a potential new and specific target in RAC1^P29S^-mutant melanoma cells. *In vitro*, CDK9 inhibition impeded the proliferation of in RAC1^P29S^-mutant melanoma cells and increased surface expression of PD-L1 and MHC Class I proteins. *In vivo*, combining CDK9 inhibition with anti-PD-1 immune checkpoint blockade significantly inhibited tumor growth only in melanomas that expressed the RAC1^P29S^ mutation. Collectively, these results establish CDK9 as a novel target in RAC1-driven melanoma that can further sensitize the tumor to anti-PD-1 immunotherapy.

## INTRODUCTION

RAC1 is a member of the RAS superfamily of small GTPases. Like other small GTPases, RAC1 functions as a molecular switch, cycling between a GTP-bound “on” state and a GDP-bound “off” state. Whole exome sequencing studies of large patient cohorts have identified 5-9% of cutaneous melanomas to harbor a *RAC1^P29S^* mutation, establishing it as the third most common hotspot mutation behind *BRAF^V600E^* and *NRAS^Q61K/R/L^* [1, 2]. The *RAC1^P29S^* mutation is a UV-induced activating mutation, resulting in preferential GTP-bound active form and sustained activation of downstream signaling [2, 3]. The *RAC1* mutation is most often found in combination with activating mutations in *BRAF* or *NRAS*, or inactivating mutations in *NF1* [4, 5]. In the case of *BRAF^V600E^*, the most common genetic driver, current first-line clinical treatments consist of combination therapy of a BRAF inhibitor and MEK inhibitor, with or without immunotherapy [6, 7]. Although *BRAF*-mutant melanomas initially respond to this combination therapy and have greatly improved clinical outcomes, adaptive resistance arises after several months of administration [6]. However, when the *RAC1^P29S^* mutation is present, melanomas display primary resistance to BRAF and MEK inhibitors [8]. The RAC1-driven mechanism of resistance remains unclear.

Compared to other cancer subtypes, melanoma has one of the highest response rates to immune checkpoint inhibition (ICI). However, predicting which patients will respond remains inaccurate, resulting in a high percentage of patient non-responders. To improve the clinical success of immunotherapy, many efforts combining ICI with adjuvant agents are currently underway. To date, there are limited studies that examine the responsiveness of tumors harboring the *RAC1^P29S^* mutation to immune therapeutic agents. Previous studies have indicated that RAC1^P29S^ expression leads to increased PD-L1 expression [9]. In addition, there are clinical data that suggest that immunotherapy is beneficial in metastatic *RAC1-mutant* melanoma [10].

This work explores the functional consequences of RAC1^P29S^ expression in melanocytes, melanoma cells, and melanoma syngeneic grafts. To characterize the RAC1^P29S^ mutation, we performed proteogenomic analysis that identified G2/M cell cycle progression as an upregulated process. Validation of our enriched pathways led to identification of CDK9 as a novel target in *RAC1^P29S^*-mutant melanoma. Furthermore, targeting CDK9 with the clinical agent dinaciclib augmented the anti-tumor immune response generated by anti-PD-1 immune checkpoint inhibition *in vivo*. These results offer a new potential treatment strategy in *RAC1*-mutant melanoma.

## MATERIALS AND METHODS

### Cell Culture

Melanocytes obtained from C57BL/6 mice were gifted by Ruth Halaban (Yale University). Melanocytes were cultured in OptiMEM supplemented with 7% horse serum, 1% FBS, 1% antibiotic, and 0.01% TPA. YUMM1.7 cells (RRID:CVCL_JK16) were purchased from ATCC and cultured in DMEM/F12 medium supplemented with 10% FBS, 1% sodium pyruvate, 1% non-essential amino acids, 1% L-glutamine, and 1% antibiotic. Cells were tested for mycoplasma every 6 weeks with MycoDect Mycoplasma Detection Kit (Alstem).

501 mel (RRID:CVCL_4633), YUROB (RRID:CVCL_B486), and YURIF (RRID:CVCL_B485) human melanoma lines were cultured in OptiMEM with 10% FBS and 1% antibiotic. IGR1 cells (RRID:CVCL_1303) were gifted from Gaudenz Danuser (University of Texas Southwestern Medical Center).

All experiments were done on cells under 15 passages. All cells were cultured at 37 degrees C, 5% CO_2_.

### Cloning

pBMN-FKBP-YFP destabilization domain plasmid was purchased from Addgene RRID:Addgene_31763. Human RAC1^P29S^ cDNA was cloned into the vector using EcoRI and XhoI restriction enzymes (New England Biolabs) and ligated using T4 ligase (New England Biolabs). Transformations were performed in Stellar competent cells (Takara). 2 μL of reaction was added to 50 μL Stellar Competent cells on ice for 30 mins followed by 45 second heat shock at 42 degrees C and resuspended in 500 μL SOC medium (Takara) and placed 1 hour shaking 200 rpm at 30 degrees C. 200 μL bacteria was then plated onto LB/AMP plate and incubated overnight at 30 degrees C. Colonies were mini-prepped with NucleoSpin Plasmid mini prep kit (Machery-Nagel). Sequences were verified by Genewiz (Azenta) plasmid sequencing services with primer sequence 5’ – TTC TGG AGC GGC GAG CAT GCA TGG AAC AGA AAC TCA TCT CTG AAG AGG - 3’. The destabilization domain was removed from the plasmid via restriction enzymes *Bam*HI and *Eco*RI (New England Biolabs) for constitutive expression in the YUMM1.7 cell lines.

### Cell Transductions

pBMN-FKBP-RAC1^P29S^ was transduced into cells through generation of a retrovirus using ϕNX cells (RRID:CVCL_H716) (purchased from Fox Chase Cancer Center Tissue Culture Facility) with Lipofectamine 2000 (Invitrogen). ϕNX were seeded 0.5 × 10^6^ cells on a 6 well cell culture plate in optiMEM (Gibco). In one Eppendorf tube, 200 μL optiMEM was mixed with 6 μL Lipofectamine 2000 Reagent. In another Eppendorf tube, 700 μL was mixed with 1 ng DNA. After 15 minutes, these tubes were mixed and incubated an additional 10 minutes. 800 μL Lipofectamine-DNA mix was added to a well with 1 mL optiMEM. Virus was collected at 24 hours and filter through a 0.45 μM filter. 4 μg/mL polybrene (Sigma-Aldrich) was added to the filtered virus, and subsequently added to target cells. Selection of positively transduced cells was enriched by sorting for HcRed positive cells on an Aria II flow cytometer (BD Biosciences).

### RAC1 Mutant Induction by Shield-1

Protein expression was induced by treatment of 1 μM Shield-1 compound (AOBIOUS) into cell culture medium for 24 hours before experiments were performed.

### Cell Proliferation

Melanocytes were seeded 100,000 cells on a 6 well plate. Cells were trypsinized with Tryp-LE express (Gibco). Cells were centrifuged and resuspended in optiMEM medium and counted on a TC 20 automated cell counter (Biorad). 3 wells were counted for each timepoint. Cells were counted every 24 hours post-seeding until 96 hours.

### Scratch Motility Assay

Cells were seeded on a density 2.5 × 10^4^ cells for well on a 12 well plate. RAC1^P29S^ was induced and cells were serum starved for 16 hours before a 20 μL pipette tip was used to scratch a vertical line across the monolayer of cells. Pictures were taken at 0 hrs and 24 hours post scratch on an EVOS fluorescent microscope on the brightfield setting. 4 wells were performed per condition and 3 pictures of each well were taken to analyze. Area was determined by ImageJ and percent cell migration was calculated using the formula: %Cell Migration = ((area_0HR_ - area_24HR_)/ area_0HR_)*100. Results were graphed in GraphPad Prism9 and statistics were calculated by a student’s t-test.

### Transwell Migration Assay

8 μM Polycarbonate membrane transwell inserts were purchased from costar and inserted into 24 well plates. Cells were seeded in serum free media in the inserts, and full serum media was placed in the well. After 24 hours, media was aspirated from the assay plate and transwell inserts. The inside of each insert was gently swabbed with a cotton round to remove non-migrated cells. Apical inserts and receiver wells were then washed in PBS and stained with crystal violet solution (6% (vol/vol) glutaraldehyde and 0.5% (wt/vol) crystal violet in H_2_0) for 10 minutes. Plates were then rinsed with tap water and visualized on a microscope. Number of stained cells was counted in 5 different views of each well (4 wells per condition). To calculate number of migrated cells the average number of cells within each field of view was divided by the microscope viewing field and then multiplied by the area of the transwell insert (0.33 cm^3^). This value was then divided by the number of cells seeded and multiplied by 100.

### Phalloidin Staining

Cells were seeded on 18 mm^2^ glass coverslip at 1 × 10^5^ inside a well on a 6 well plate. Cells were washed in PBS and fixed in 4% paraformaldehyde (Electron Microscopy Sciences, 15710) in the dark for 15 minutes. Slides were wash 2 times in PBS followed by permeabilization with 0.1% Triton X-100 in PBS for 15 minutes. Slides were washed 2 times in PBS and incubated in Alexa Fluor 488 Phalloidin (ThermoFisher Scientific, A12379) at 1:400 diluted in PBS with 1% BSA. Cells were mounted in ProLong Gold Antifade Mountant with DAPI (Thermo Fisher Scientific) and allowed to dry overnight. Images were acquired on a laser scanning confocal microscope (Leica TSC SP8 Confocal).

### CRISPR-mediated CDK9 Knockout and Cell Electroporation

Cas9 nuclease alone or Cas9 nuclease and sgCDK9 were electroporated into YUMM1.7 cells at a ratio of 9:1. sgCDK9 pooled sequences: (C*U*C*CCUUCAGAUCCUGCACA), (U*G*G*GCUGGCUAUUCUUAGCU), (C*G*A*GCAGCAGCUCCGGAGGC). Cas9 nuclease and sgCDK9 pool were purchased from Synthego. RNP complexes were assembled by mixing 30 μM CDK9 sgRNA with 20 μM Cas9 and 3.5 Resuspension buffer (Invitrogen). RNP mix was incubated at room temperature for 10 minutes before electroporation. 1 × 10^6^ YUMM1.7 cells were resuspended in 50 μL resuspension buffer R and cell-RNP mixture was produced by mixing 5 μL cell suspension with 7 μL RNP mixture. 10 μL cell-RNP mixture was aspirated into a 10 μL Neon tip and electroporated by a Neon Transfection System (Invitrogen). YUMM1.7 cells were electroporated at 1500 V, 20 ms width, for 2 pulses.

### RNA Sequencing

Total RNA was isolated with TRI-reagent (Invitrogen) and chloroform (Sigma-Aldrich) per manufacturer’s instructions. RNA was sequenced by the Next Generation Sequencing Facilities at Fox Chase Cancer Center. Samples were sequenced on an Illumina HiSeq 2500 sequencer. Raw data was analyzed by Fox Chase Cancer Center Biostatistics and Bioinformatics Facility.

### Multiplex Inhibitor Beads and Mass Spectrometry

MIBs/MS was performed in collaboration with the James Duncan lab at Fox Chase Cancer Center.

#### Lysate Preparation

Cells were scraped from 150 mm plates with MIBs Lysis buffer (50mM HEPES pH 7.5, 150 mM NaCl, 0.5% Triton X-100, 1 mM EDTA, 1 mM EGTA, 10 mM NaF, 2.5 mM NaVO4, Protease inhibitor cocktail (Roche), phosphatase inhibitor cocktail 2 (Sigma P5726), Phosphatase inhibitor cocktail 3 (Sigma-P0044)) and added to a 15 mL conical tube. Lysates were sonicated using a tip sonicator at 3 x 30 pulses at 20% duty cycle. Lysates were then transferred to Eppendorf tubes and centrifuged at 4 degrees for 15 minutes at max speed. Supernatants were then filtered through 0.45 μm CA membrane filter. Protein concertation was determined by BCA assay. 5 mg of protein was used for each replicate for subsequent MIBs/MS steps (n=3 per condition).

#### MIBs/MS

Kinases were isolated by flowing lysates over kinase inhibitor-conjugated Sepharose beads (purvalanol B, VI16832, PP58 and CTx-0294885 beads) in 10 mL gravity flow columns. Columns were washed 2 times in high salt buffer (50mM HEPES pH 7.5, 1 M NaCl, 0.5% Triton X-100, 1 mM EDTA, 1 mM EGTA) followed by 1 time in low salt buffer (50 mM HEPES pH 7.5, 150 mM NaCl, 0.5% Triton X-100, 1 mM EDTA, 1 mM EGTA). Kinases were eluted from the column with elution buffer (0.5% SDS, 1% β-mercaptoethanol in 0.1 M Tris-HCl pH 6.8) and boiled. Eluted peptides were incubated with 5 mM DTT at 65 degrees C followed by 20 mM iodoacetamide at room temp for reduction and alkylation. Alkylation was quenched with DTT. Samples were then concentrated to 100 μL with Millipore 10kD cutoff spin concentrators. Detergent was removed by chloroform/methanol extraction, and protein pellet was resuspended in 50 mM ammonium bicarbonate and digested with sequencing grade trypsin (Promega) overnight at 37 degrees C. Digested peptides were then cleaned with PepClean C18 spin columns (ThermoFisher Scientific) and dried in a speed-vacuum and resuspended in 50 μL 0.1% formic acid and extracted by ethyl acetate. Any additional ethyl acetate was allowed to evaporate, and peptides were dried in a speed vacuum, followed by LC-/MS/MS analysis.

### Cell Cycle Profile Analysis

Cells were trypsinized in TrpLE-express (Gibco) and fixed dropwise while vortexing in ice cold 70% ethanol and stored at −20°C. Cells were then stained with 1 mg/mL propidium iodide (Invitrogen) with 1 mg/mL RNase A (Sigma Aldrich) in PBS. Cell cycle was measured on a FACSymphony A5 cytometer (BD Biosciences). Data was analyzed in FlowJo FlowJo (RRID:SCR_008520) and graphed on GraphPad Prism 9.

### H2B-GFP Live Cell Imaging

Cells were transduced with an H2B-GFP plasmid (Addgene 11680) and were seeded 0.25 × 10^5^ cells onto 6 well plates and treated with +/-SHLD 1 ligand. Brightfield and GFP photos were taken every 15 minutes for 24 hours to track the mitotic cell cycle phase. Images were analyzed on ImageJ and mitosis was quantified by onset of mitosis (visible condensation of chromosomes) to anaphase (clear separation of sister chromatids).

### Pharmacologic Inhibitor Treatments and Viability Assays

Cells were seeded from 750-2000 cells per well on a white 96 well plated and allowed to adhere overnight. 24 hours after transgene induction (when necessary), cells were treated with a dose titration of pharmacologic inhibitors. After an incubation time of 48 hours, viability was measured by Cell Titer Glo 2.0 assay (Promega) per manufacturer’s protocol. Plates were read on a luminescent plate reader (Perkin Elmer Envision Plate Reader). Drug treatment dose ranges included Alisertib: 0.0001-50 μM, Volasertib: 0.0001-50 μM, Tozasertib: 0.0001-50 μM, Dinaciclib: 1-100 nM, R03306: 0.1-50 μM, NU2058: 0.1-100 μM, SU9516: 0.1-75 μM, Abemaceclib: 0.01-325 μM, AT519: 0.1-4 μM, NVP2: 0.1-1000 nM, JQ-1: 1-1000 nM.

### Immunoblot

Protein was isolated using RIPA buffer with protease (Sigma Aldrich) and phosphatase cocktail inhibitors (Sigma Aldrich). Cells were washed twice with 1X PBS on ice and incubated for 5 minutes in RIPA buffer. Cells were then scraped off and sonicated for 3 pulses at 20% (550 sonic dismembrator, Fisher Scientific). Cells were then centrifuged for 15 minutes at 15,000 x g at 4 degrees. The supernatant was transferred to a new Eppendorf tube and protein was quantified with Quick Start Bradford 1X Dye Reagent (BioRad). Samples were added to 4X Laemmli Sample Buffer (BioRad) plus 10% β-mercaptoethanol and boiled for 5 minutes. 20 μg of protein was loaded on a Mini-PROTEAN TGX gel 4-20% gradient pre-cast gels (BioRad). Protein was transferred to 0.45 μm nitrocellulose membrane (Biorad) and blocked per manufacturer’s instructions in either 5% milk-TBST or 5% BSA-TBST. Antibody dilutions were prepared in 5% BSA-TBST and incubated overnight with gentle agitation at the following dilutions phospho-AKT T308 1:2000 (Cell Signaling Technology Cat# 13038, RRID:AB_2629447), AKT 1:1000 (Cell Signaling Technology Cat# 9272, RRID:AB_329827), phospho-ERK 1/2 Y202/204 1:2000 (Cell Signaling Technology Cat# 4370, RRID:AB_2315112), ERK 1:1000 (Cell Signaling Technology Cat# 4695, RRID:AB_390779), phospho-PAK 1/2 S199/204, S192/197 1:1000 (Cell Signaling Technology Cat# 2605, RRID:AB_2160222), PAK 1/2/3 1:1000 (Cell Signaling Technology Cat# 2604, RRID:AB_2160225), Myc-Tag 1:1000 (Cell Signaling Technology Cat# 2276, RRID:AB_331783), GAPDH 1:1000 (Cell Signaling Technology Cat# 2118, RRID:AB_561053), Vinculin 1:1000 (Cell Signaling Technology Cat# 13901, RRID:AB_2728768), CDK9 1:1000 (Cell Signaling Technology Cat# 2316, RRID:AB_2291505), phospho-RNA Polymerase II S2 1:1000 (Millipore Cat# 04-1571, RRID:AB_11212363), PTEN 1:1000 (Cell Signaling Technology Cat# 9552, RRID:AB_10694066).

Secondary antibodies were Peroxidase AffiniPure Goat Anti-Mouse IgG (H+L) 1:10000 (Jackson ImmunoResearch Labs Cat# 115-035-003, RRID:AB_10015289) and Peroxidase AffiniPure Goat Anti-Rabbit IgG (H+L) 1:10000 (Jackson ImmunoResearch Labs Cat# 111-035-003, RRID:AB_2313567).

### Live/Dead Assays

Cells were suspended by trypsinization with Tryp-LE Express (Gibco). Cells were stained using LIVE/DEAD Viability/Cytotoxicity Kit (Invitrogen, L3224) per manufacturer’s protocol. Cells were analyzed via flow cytometry on a Symphony A5 cytometer (BD Biosciences). Compensation controls were used to gate for live and dead cells on PE and BB630 fluorescent channels. Data analysis was performed on FlowJo and graphed on GraphPad Prism 9.

### Crystal Violet Staining

Crystal violet solution fixation solution consisted of 6% (vol/vol) glutaraldehyde and 0.5% (wt/vol) crystal violet in H_2_0. Solution was added to cells in a 6 well plate and incubated for 30 mins. Cells were then rinsed in tap water and allowed to dry.

### Cell Surface Staining by Flow Cytometry

Cells were suspended using Tryp-LE Express (Gibco). Cells were fixed in 4% PFA (Electron Microscopy Sciences) for 15 minutes and washed 3 times in PBS before staining. Cells were stained with fluorescent-conjugated primary antibodies rotating overnight at 4 degrees Celsius. Cells were then washed 3 times in PBS and analyzed by flow cytometry using a Symphony A5 cytometer (BD Biosciences). Data was analyzed with FlowJo software and graphed on GraphPad Prism 9. Statistics were calculated by GraphPad Prism.

Antibodies used: PD-L1 0.125 μg/test (Thermo Fisher Scientific Cat# 53-5982-82, RRID:AB_2811871), MHC Class I 1 μg/test (Thermo Fisher Scientific Cat# 11-5958-82, RRID:AB_11149502), IgG Isotype Control 0.125 μg/test (Thermo Fisher Scientific Cat# 11-4888-81, RRID:AB_470037).

### *In vivo* Experiments

6-week-old male C57Bl/6J mice were purchased from Jackson Laboratories (RRID:IMSR_JAX:000664) and allowed to acclimate for 2 weeks. Male mice were used in this study as the YUMM1.7 cell line originated from a male mouse, and female hosts can reject grafts due to antigens from the Y-chromosome. Post acclimatization, syngeneic grafts were established by subcutaneous injections of 1,000,000 cells in PBS. Cells injected were either the YUMM1.7 cell line or YUMM1.7 + RAC1^P29S^. Tumor sizes were monitored daily until tumors reached 100 mm^3^. Tumors were measured with vernier calipers and tumor volume was calculated by ((length x width^2^) x 0.5). Mice were randomly divided into 4 cohorts respective of tumor genotype: Vehicle, dinaciclib, anti-PD1 antibody, or dinaciclib + anti-PD1 antibody. Vehicle consisted of 10mg/mL rat IgG2a isotype control diluted in *InVivo*Pure 6.5 dilution buffer and 20% hydroxypropyl-β-cyclodextrin. Dinaciclib was dissolved in 20% hydroxypropyl-β-cyclodextrin and dosed at 30 mg/kg via intraperitoneal injection every three days. Anti-mouse PD-1 (RMP1-14) was diluted in *InVivo*Pure pH 7.0 dilution buffer and dosed 10mg/kg every 4 days via intraperitoneal injection. Tumors and weight were measured every three days post regimen start for 21 days. Humane endpoints included tumor mass over 10% of body weight, tumor length greater than 20 mm, or tumor ulceration. Dinaciclib (SCH727965) was purchased from Selleck Chemicals (S2768). Dinaciclib was diluted in 20% hydroxypropyl-β-cyclodextrin (w/v) purchased from Sigma-Aldrich (H107-5G). The following reagents were purchased from BioXCell: *InVivo*MAb anti-mouse PD-1 (Clone RMP1-14) (BE0146) which was diluted in *InVivo*Pure pH 7.0 dilution buffer (IP0070). *InVivo*MAb rat IgG2a isotype control, anti-trinitrophenol (Clone 2A3) (BE0089) and was diluted in *InVivo*Pure pH 6.5 dilution buffer (IP0065).

### Study Approval

All animal experiments were performed according to the approved protocols of the Fox Chase Cancer Center Institutional Animal Care and Use Committee.

### Immunohistochemical (IHC) staining

Mouse tumor tissues were collected and fixed in 10% neutral buffered formalin for 24-48 hrs, dehydrated and embedded in paraffin. Hematoxylin and eosin (H&E) stained sections were used for morphological evaluation purposes and 5-μm unstained sections for immunohistochemical (IHC) studies.

Immunohistochemical staining was performed on a VENTANA Discovery XT automated staining instrument (Ventana Medical Systems) using VENTANA reagents according to the manufacturer’s instructions. Briefly, slides were de-paraffinized using EZ Prep solution (cat # 950–102) for 16 min at 72 °C. Epitope retrieval was accomplished with CC1 solution (EDTA, ph 9.0. cat # 950–224) at high temperature (eg, 95–100 °C) for 32 min. Rabbit primary antibodies (anti-mouse CD8, 1:50, (Cell Signaling Technology Cat# 98941, RRID:AB_2756376); CD11C: 1:100 (Cell Signaling Technology Cat# 39143, RRID:AB_2924836); CD86 (Cell Signaling Technology Cat# 19589, RRID:AB_2892094), tittered with a TBS antibody diluent into user fillable dispensers for use on the automated stainer. Immune complex was detected using the Ventana OmniMap anti-Rabbit detection kit (760-4311) and developed using the VENTANA ChromMap DAB detection kit (cat # 760-159) according to the manufacturer’s instructions. Slides were then counterstained with hematoxylin II (cat # 790-2208) for 8 min, followed by Bluing reagent (cat # 760-2037) for 4 min. The slides were then dehydrated with ethanol series, cleared in xylene, and mounted. As a negative control, the primary antibody was replaced with normal rabbit IgG to confirm absence of specific staining.

### Quantitative Image analysis

Immunostained slides were scanned using an Aperio ScanScope CS 5 slide scanner (Aperio, Vista, CA, USA). Scanned images were then viewed with Aperio’s image viewer software (ImageScope, version 11.1.2.760, Aperio). Selected regions of interest were outlined manually by a pathologist. The immune cells quantifications were performed with Aperio V9 or positive pixel count (PPC) algorithm.

### Statistical Analysis

Statistical Analysis was done in GraphPad Prism (RRID:SCR_002798). For comparisons of two groups, statistical analysis was performed using Student’s t-test: * *P* < 0.05; ** *P* < 0.01; *** *P* < 0.001; **** *P* < 0.0001. Immunoblot results are representative images of three experiments and quantifications were analyzed by ImageJ (RRID:SCR_003070). Analysis of multiple groups was performed using ordinary two-way ANOVA, Tukey’s multiple comparisons test.

### Data Availability Statement

All other raw data generated in this study are available from the corresponding author upon request.

## RESULTS

### Characterization of *RAC1^P29S^* oncogenic properties

To characterize the RAC1^P29S^ effect on downstream signaling, morphology, and cell proliferation, we generated an inducible cell system of RAC1^P29S^ expression. Melanocytes isolated from C57BL/6 mice were obtained and genetically altered via CRISPR/Cas technology to knock out *Pten*, since this is a commonly altered tumor suppressor in cutaneous melanoma and mouse models of combined *Braf*/*Pten* mutations have been well-studied [4, 5] (Supplemental Figure 1). A plasmid containing a destabilization domain (DD) fused to *RAC1^P29S^* was then introduced into these C57BL/6 *Pten^-/-^* melanocytes (Figure 1A). Under basal conditions, the DD results in translation of an unstable protein, which is immediately targeted and degraded by the proteasome. However, upon treatment with Shield1 ligand (SHLD1), the DD is stabilized, and the recombinant protein is expressed [11]. This expression system was originally produced in a non-transformed C57BL/6 cell line to delineate molecular consequences specific to the RAC1^P29S^ mutation without confounding signaling inputs from other potential oncogenes. After establishing cellular RAC1^P29S^ expression, we examined the RAC1^P29S^ effect on downstream MAPK and AKT signaling, both of which lead to cell proliferation and survival. RAC1^P29S^ resulted in a slight increase in phospho-PAK, phospho-ERK and phospho-AKT T308 (Figure 1B). Melanocytes that expressed RAC1^P29S^ also displayed faster proliferation rates than non P29S-expressing counterparts (Figure 1C). As RAC1 is a master regulator of actin dynamics, we explored the effect on motility, migration, and actin structure. To further examine the effects of RAC1^P29S^ on cell motility and migration, we performed scratch and transwell assays. RAC1^P29S^-expressing melanocytes exhibited higher motility, as a significantly higher number of cells migrated into the scratch area (Figure 1D). Additionally, we observed higher cell migration in a transwell assay (Figure 1E). Phalloidin staining of actin displayed increased membrane ruffling and lamellipodia formation compared to melanocytes that did not express the P29S mutation (Figure 1F). This data concludes that RAC1^P29S^ is an activating mutation that leads to MAPK activation, lamellipodia formation, and increased motility and migration. However, as these effects are modest, there may be additional mechanisms that contribute to RAC1-driven oncogenicity.

**Figure 1.**
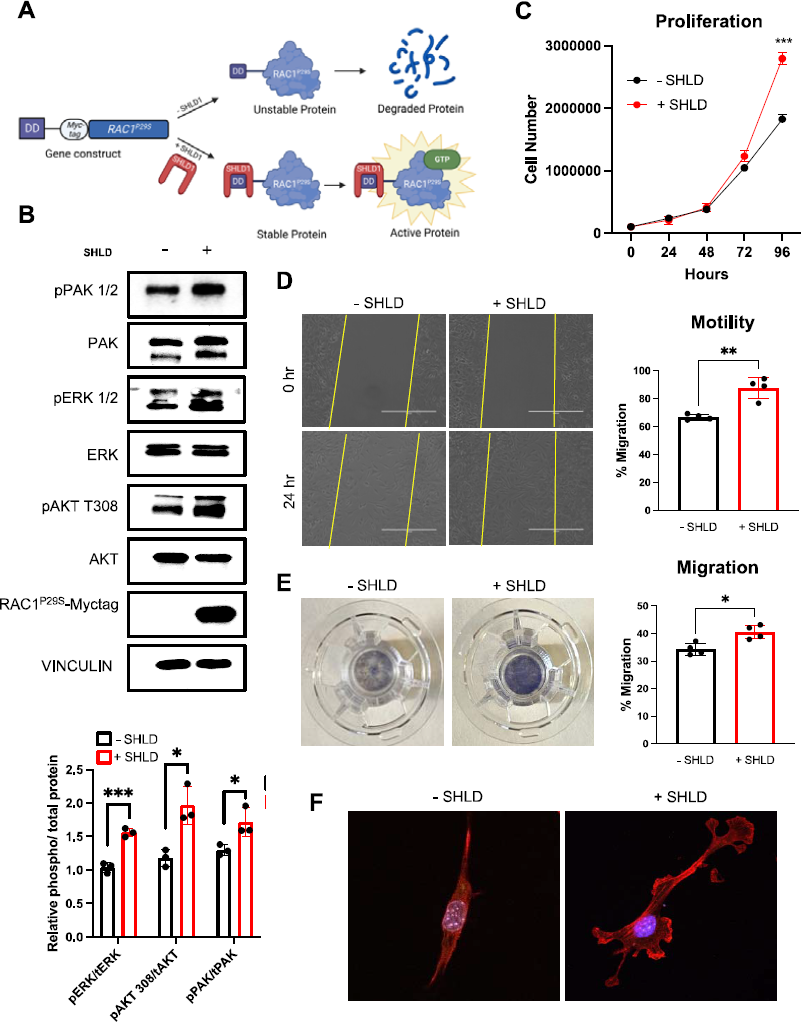
Characterization of *RAC1P29S* oncogenic properties. **A.** Schematic of the Shield 1 (SHLD1) regulated gene expression. SHLD1 bind to the destabilization domain on translated protein and prevents it from being degraded. Active protein proceeds to propagate its downstream cellular effects. Without SHLD1, the protein is immediately degraded. **B.** Downstream signaling of pPAK1(S199/204)/PAK2 (S192/197), pERK1/2 (T202/T204), and pAKT (T308) and phospho-total protein quantification from 3 repeated blots. Bars on graphs represent means with standard deviation. **C.** Cell proliferation measured every 24 hours. Graph represents the mean value and standard deviation (n=3). **D.** Representative images of motility scratch assay at 0 and 24 hours. Initial scratch area is highlighted in yellow. Graph represents the mean value and standard deviation (n=4). **E.** Representative transwell inserts and quantified migration after 24 hours (n=4). **F.** Representative images of phalloidin immunofluorescent stain of cells.

### Proteogenomic analysis of RAC1^P29S^ melanocytic expression reveals upregulation of cell cycle processes

To determine the molecular changes induced by RAC1^P29S^ expression, we performed RNA-sequencing (RNA-seq) coupled with multiplexed inhibitor bead-mass spectrometry (MIBs/MS). RNA-seq analysis of expressed recombinant RAC1^P29S^ mouse-derived melanocytes resulted in 600 differentially expressed genes. E2F_TARGETS and G2M_CHECKPOINT were the top two enriched hallmark gene sets (Figure 2A, B). Other cell cycle related processes included MYC_TARGETS_V1 and MITOTIC_SPINDLE (Figure 2A). The top 10 enriched REACTOME gene sets were all cell cycle events, indicating RAC1^P29S^ has a role in influencing cell cycle progression, consistent with previous studies on RAC1 (Figure 2C) [12, 13]. In addition, INFLAMMATORY_RESPONSE and IL-2_STAT5_SIGNALING hallmark gene sets were enriched, suggesting a potential function in modulating inflammation (Figure 2A). Ingenuity Pathway Analysis (IPA) identified the top altered upstream regulators in response to RAC1^P29S^ expression (Figure 2D). Top activated upstream regulators included mainly transcription factors that have been reported to promote expression of cell cycle regulating genes such as E2F proteins, YAP1, CCND1, and FOXM1. Accordingly, the top inactive upstream regulators were tumor suppressors and cell cycle inhibitors including COPS5, SMARCB1, CDKN2A, and RB1 (Figure 2D). Interestingly, the activity of several transcriptional regulators was altered as well, including MED1 (a subunit of the mediator complex which helps position RNA polymerase II on DNA promoter regions), HDAC1 (histone deacetylase), and PRDM16 (histone methyltransferase) (Figure 2D). MIBs/MS further implicated cell cycle progression as an enriched pathway in RAC1^P29S^-expressing cells through identification of 7 kinases involved specifically in G2/M transition (Figure 2E).

**Figure 2.**
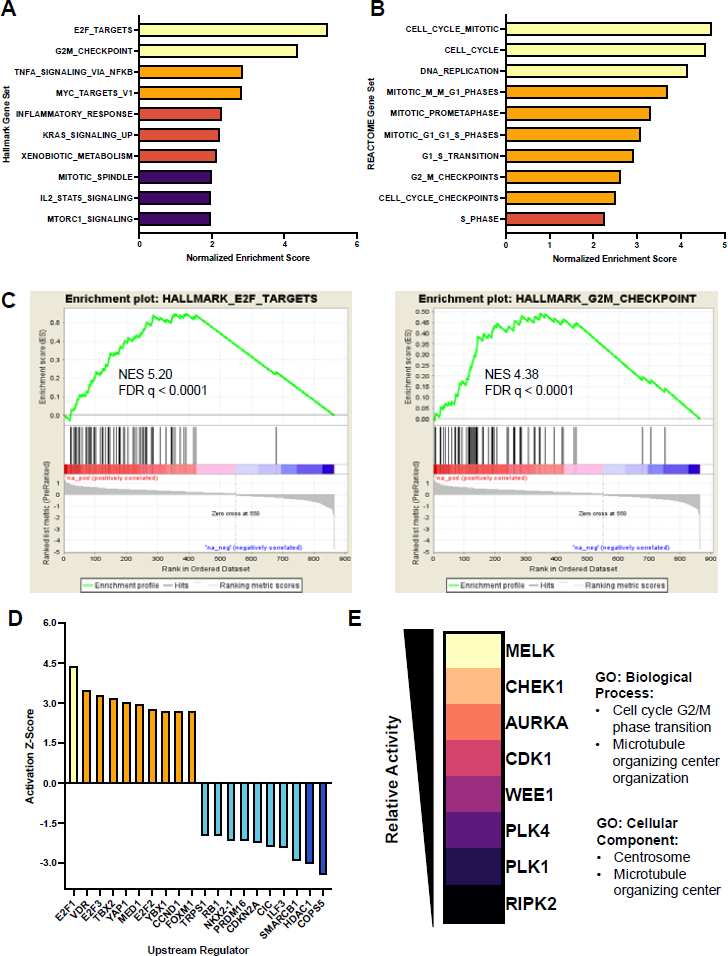
Proteogenomic analysis of RAC1^P29S^ melanocyte expression reveals upregulation of cell cycle processes. **A.** Normalized enrichment scores (NES) of Hallmark gene sets Molecular signatures Database (MSigDB). NES > 3.5 (yellow), NES > 2.5 (orange), NES >2 (peach), NES >1.5 (purple). FDR < 0.05 cut off. **B.** NES of REACTOME gene sets (MsigDB). NES > 3.5 (yellow), NES > 2.5 (orange), NES >2 (peach). FDR < 0.05 cut off. **C.** GSEA plots of top two enriched hallmark gene sets. **D.** Activation Z-Score of top active/inactive upstream regulators determined by QIAGEN Ingenuity Pathway Analysis (IPA). Z-score >4 (red), Z-score > 2 (orage), Z score > −2.5 (cyan), Z-score > −4 (blue). **E.** Enriched kinases identified by MIBs/MS and associated Gene Ontology. Kinases listed as most active (yellow) to least active (black). Gene Ontology corresponds to enriched MIBs kinases.

To further investigate the RAC1^P29S^ influence on the cell cycle, the cell cycle profile was examined by flow cytometry. RAC1^P29S^-expressing melanocytes showed a lower percentage of cells in G2/M phase, indicating the RAC1^P29S^ mutation may lead to accelerated transition through the G2/M cell cycle checkpoint (Supplemental Figure 1A). P29S-expressing melanocytes also exhibited a higher percentage of cells in S phase, indicating increased DNA replication, consistent with gene set enrichment analysis (GSEA) (Supplemental Figure 1A). Additionally, live H2B-GFP cell tracking resulted in less time spent in the prophase to anaphase stages of mitosis in the RAC1-mutant cells, further suggesting a role in mitotic cell cycle progression (Supplemental Figure 1B).

### RAC1^P29S^-expressing melanocytes show increased sensitivity to CDK9 inhibition

Due to the strong G2/M cell cycle phase transition signature yielded by the proteogenomic and cell cycle analysis, we next further examined the kinase signaling cascade that promotes G2 to M phase transition to investigate potential targeting vulnerabilities of RAC1^P29S^-expressing melanocytes. PAK1 phosphorylates Aurora kinases (AURK) and Polo-like kinase 1 (PLK1), which in turn phosphorylates CDC25, ultimately leading to CDK1 phosphorylation, activation, and G2/M phase transition [14, 15] (Figure 3A). While there were no differential sensitivities to alisertib (AURKA inhibitor), tozasertib (pan-AURK inhibitor), or volasertib (PLK1 inhibitor), melanocytes expressing a RAC1^P29S^ mutation were significantly more sensitive to dinaciclib (CDK1,2,5,9 inhibitor) (Figure 3B). However, as dinaciclib is not specific to CDK1, we performed a CDK inhibitor viability panel to determine which CDK target conferred the observed sensitivity (Figure 3C). Interestingly, the compounds that targeted CDK9 were all significantly more cytotoxic to melanocytes expressing the RAC1^P29S^ mutation (Figure 3C,D).

**Figure 3.**
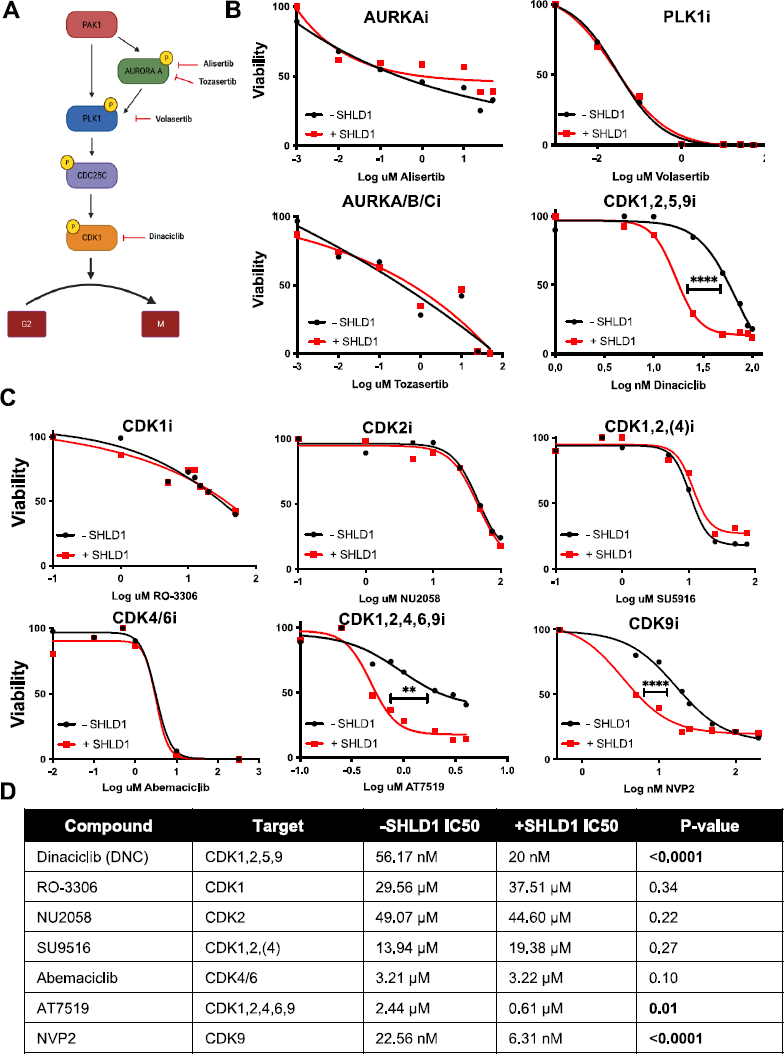
RAC1^P29S^ expression confers increased sensitivity to CDK9 inhibitors. **A.** Simplified signaling cascade leading to G2/M cell cycle transition. **B.** Cell viability in response to alisertib, volasertib, tozasertib, and dinaciclib. p < 0.0001 denoted by ****. **C.** Cell viability in response to various CDK inhibitors. p < 0.0001 denoted by ****, p < 0.001 denoted by ***, p < 0.01 denoted by **. p < 0.05 denoted by *. **D.** Compound, target and IC50 values. Values calculated from nonlinear regression dose response curve, determined by GraphPad Prism 9.

To further examine RAC1^P29S^-mediated sensitivity to CDK9, we generated a constitutive RAC1^P29S^-expressing melanoma cell line. YUMM1.7 cell lines (*Braf^V600E^, Pten^null^, Cdkn2d^null^)* were transfected with myc-tagged *RAC1^P29S^* (Figure 4A). YUMM1.7 cells that expressed RAC1^P29S^ were significantly more sensitive to both dinaciclib and the CDK9 specific inhibitor NVP2 (Figure 4B,C). To confirm this sensitivity was indeed due to CDK9 inhibition we knocked out CDK9 using CRISPR-Cas9 (Figure 4D). Either Cas9 plus sgCDK9 or Cas9 alone were electroporated into YUMM1.7 or YUMM1.7 plus RAC1^P29S^ cells. 48 hours post electroporation, YUMM1.7 cells expressing RAC1^P29S^ had a significantly higher percent of dead cells compared to those that did not express mutant RAC1 (Figure 4E). After 120 hours, crystal violet staining showed RAC1^P29S^ expressing YUMM1.7 melanoma cells were no longer viable, further implicating CDK9 as a vital protein in RAC1^P29S^-expressing cells (Figure 4F).

**Figure 4.**
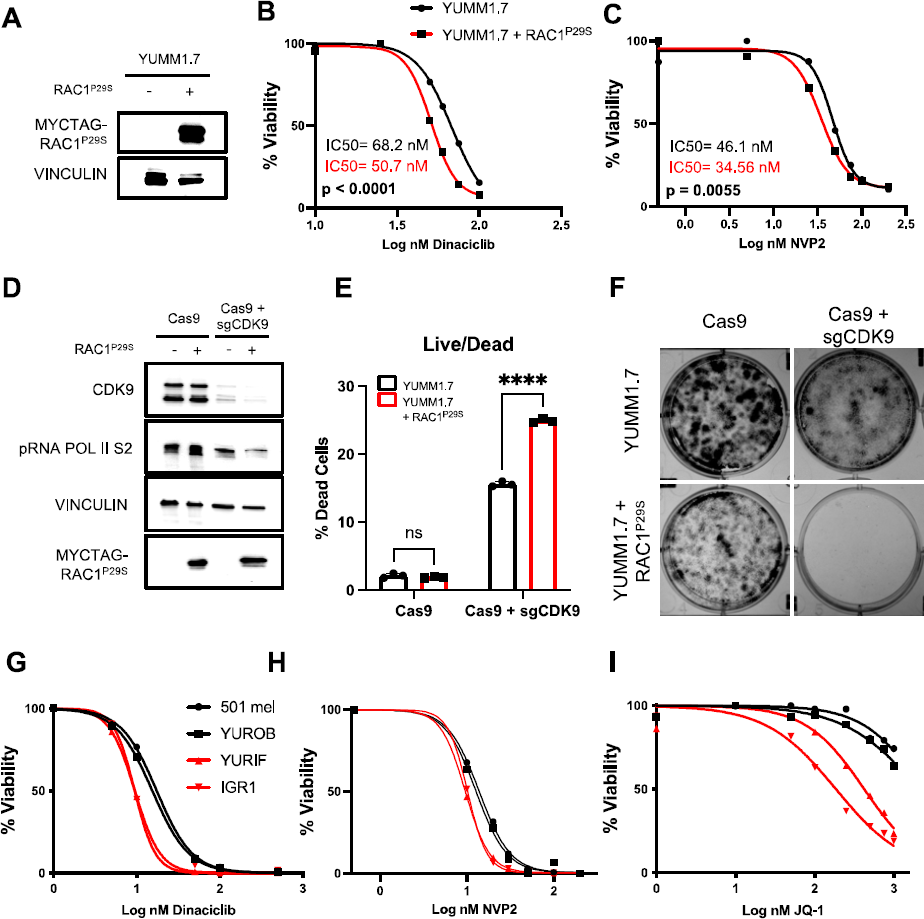
RAC1^P29S^ expression confers increased sensitivity to CDK9 inhibition and genetic knockout in melanoma cells. **A.** Immunoblot of YUMM1.7 or YUMM1.7 with constitutive expression of myc-tagged RAC1^P29S^. **B.** Viability curves after dose titrations of dinaciclib (10 nM – 100 nM) or **C.** NVP2 (0.5 nM – 200 nM). Nonlinear regression curves fit and p value calculated in GraphPad Prism 9. Drugs were incubated with cells for 48 hours. **C.** CRISPR mediated knockout of YUMM1.7 melanoma cells +/-RAC1^P29S^. Cells were electroporated with Cas9 alone or Cas9 + sgCDK9. **D.** Quantification of LIVE/DEAD assay 48 hours post electroporation. Samples were analyzed via flow cytometry. p < 0.0001 **E.** Crystal violet staining 120 hours post electroporation. **F.** Viability curves after dose titrations of dinaciclib (1 nM – 500 nM), **H.** NVP2 (0.5 nM – 200nM), and **I.** JQ-1 (1 nM – 1000 nM) incubated for 48 hours. Nonlinear dose response curves generated by GraphPad Prism 9. P values calculated by Tukey’s multiple comparisons test, 2way ANOVA. P < 0.0001 (****).

CDK9 is a non-canonical CDK, which has a primary function in transcriptional regulation through phosphorylation of Serine 2 on the C terminal domain (CTD) tail of RNA polymerase II [16, 17]. In cancer, aberrant activity of CDK9 promotes transcription of genes that facilitate cell survival, such as those that promote cell cycle progression [18]. To confirm that sensitivity of cells harboring a *RAC1^P29S^* mutation is transferrable across multiple melanoma cell lines and genetic backgrounds, we examined CDK9 sensitivity across several human melanoma lines, including those that harbor endogenous *RAC1^P29S^* mutations. The melanoma cells bearing P29S mutations were significantly more sensitive to dinaciclib and NVP2 (Figure 4G-H).

We also tested the effects of a BRD4 inhibitor, as active CDK9 and Cyclin T form the positive transcription elongation factor b (P-TEFb) complex, which associates with bromodomain-containing protein 4 (BRD4) to activate RNA polymerase II transcriptional elongation [19]. We found that human melanoma lines harboring a *RAC1^P29S^* mutation are also more sensitive to inhibition of BRD4 with JQ-1 (Figure 4I).

Although CDK9 does not primarily regulate the cell cycle as a classic CDK, previous studies have linked CDK9 with promotion of transcription of G2/M genes [18].

Accordingly, melanocytes trend toward G2/M arrest after CDK9 inhibition with dinaciclib or NVP2 for 16 hours, with a lower dose necessary for G2/M arrest in RAC1^P29S^-expressing cells (Supplemental Figure 2A-B).

These data suggest that RAC1^P29S^ induces molecular changes that lead to increased dependency on CDK9, presenting a potential therapeutic target in this subset of melanoma. Additionally, the cell cycle and G2/M signature seen in the RNA-seq and MIBs/MS datasets may be influenced by CDK9 modulation of RNA polymerase II transcribed gene regulation.

### CDK9 inhibition increase surface expression of PD-L1 and Class I MHC in RAC1 mutant melanocytes

Several recent studies have reported pharmacologic inhibition of CDKs to regulate PD-L1 expression and sensitize the tumor microenvironment to immunotherapy [20–26]. Given the unique sensitivity of RAC1^P29S^ mutant melanoma cells to CDK9 inhibition, we next explored the consequences of CDK9 inhibition on cellular immunogenicity. Dinaciclib and NVP2 both decreased phosphorylated RNA polymerase II at serine 2 (Figure 5A,D). Following dinaciclib treatment, both surface expression of PD-L1 and major histocompatibility complex (MHC) Class I are significantly more elevated in P29S-expressing melanocytes (Figure 5B). Cells treated with NVP2 also exhibited an increase in surface PD-L1 and MHC Class I expression (Figure 5C). The same increase in PD-L1 and MHC Class I was also seen upon dinaciclib treatment in YUMM1.7 plus RAC1^P29S^ melanoma cell lines (Figure 5E) and upon NVP2 treatment (Figure 5F).

**Figure 5.**
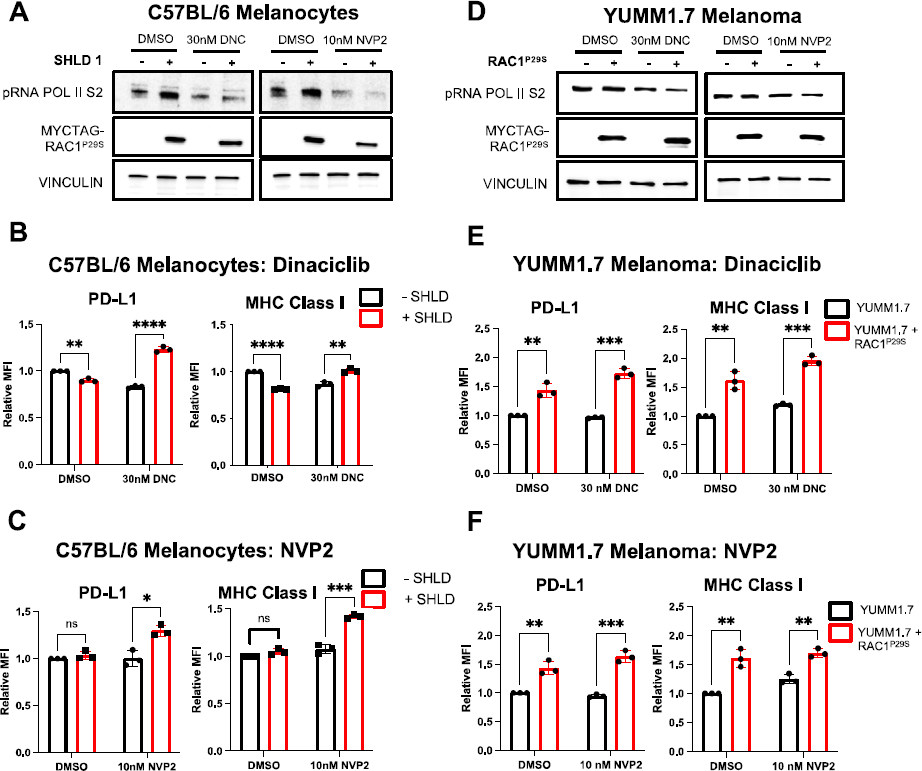
CDK9 inhibition increase surface expression of PD-L1 and Class I MHC in RAC1 mutant melanocytes. **A.** Representative immunoblot of phosphorylated RNA polymerase II Ser2 after pharmacological inhibition by dinaciclib (DNC) or NVP2 in C57BL/6 melanocytes. **B.** Surface expression of PD-L1 or MHC Class I 72 hours after DNC treatment in C57Bl/6 based cell lines +/-SHLD. PD-L1: DMSO p = 0.001. 30 nM DNC p = 0.00003. MHC Class I: DMSO p <0.001. 30 nM DNC p = 0.002. All values normalized to DMSO-SHLD. **C.** Surface expression of PD-L1 or MHC Class I 72 hours after NVP2 treatment in C57Bl/6 based cell lines. PD-L1: DMSO p = 0.203. 30 nM DNC p = 0.007. MHC Class I: p = 0.057. 30 nM DNC p = 0.0003. All values normalized to DMSO-SHLD. **D.** Representative immunoblot of phosphorylated RNA polymerase II Ser2 after pharmacological inhibition by dinaciclib (DNC) or NVP2 in YUMM1.7 melanoma cell lines. **E.** Surface expression of PD-L1 or MHC Class I 72 hours after DNC treatment in YUMM1.7 based cell lines. PD-L1: DMSO p = 0.004. 30 nM DNC p = 0.0001. MHC Class I: p = 0.003. 30 nM DNC p = 0.0001. All values normalized to YUMM1.7 DMSO. **F.** Surface expression of PD-L1 or MHC Class I 72 hours after NVP2 treatment in YUMM1.7 based cell lines. PD-L1: DMSO p = 0.004. 30 nM DNC p = 0.0003. MHC Class I: p = 0.003. 30 nM DNC p = 0.002. All values normalized to YUMM1.7 DMSO.

MHC class I is found on the cell surface of all nucleated cells and plays an instrumental role in antigen presentation and T-cell activation. Upregulation of MHC Class I could lead to more antigen presentation and a more robust anti-tumor immune response.

These results indicate CDK9 inhibition could increase the immunogenicity of the tumor microenvironment and potentially enhance immune checkpoint inhibition response.

### Combination therapy of CDK9 inhibition and anti-PD-1 immunotherapy significantly decreases tumor growth *in vivo*

As our *in vitro* data indicated CDK9 inhibition as a potential enhancer of immune checkpoint inhibition, we next tested the efficacy of CDK9 inhibition with anti-PD-1 ICI in a syngeneic murine model. Subcutaneous (s.c.) tumors were established in immune competent C57BL/6 mice with YUMM1.7 (*Braf^V600E^, Pten^null^, Cdkn2d^null^)* melanoma cells, or YUMM1.7 cells constitutively expressing the RAC1^P29S^ mutation. Once tumors reached approximately 100 mm^3^, mice were randomly allocated to 4 treatment groups and either treated with vehicle (20% (2-Hydroxypropyl)-β-cyclodextrin) + IgG isotype control, 30 mg/kg dinaciclib, 10 mg/kg anti-PD-1 Ab, or a combination of dinaciclib plus anti-PD-1 Ab. Dinaciclib was treated every 3 days, while anti-PD-1 Ab was administered every 4 days (Figure 6A). YUMM1.7 tumors did not significantly respond to either monotherapies or the combination therapy (Figure 6B). Tumor growth inhibition (TGI) of dinaciclib alone was 1%, anti-PD1 antibody alone was 0%, and combination treatment was 16%. Contrastingly, YUMM1.7 tumors that expressed RAC1^P29S^ were significantly repressed with treatment of either monotherapies and overall inhibited by combination treatment (Figure 6C). Dinaciclib alone had a TGI of 72%, anti-PD-1 Ab alone had a TGI of 66%, and dinaciclib and anti-PD-1 Ab combination had a TGI of 97%, with one complete responder. YUMM1.7 tumor response greatly varied to dinaciclib and anti-PD1 Ab combination, while YUMM1.7 + RAC1^P29S^ tumors were all suppressed, with 4/5 tumors smaller than their size at the beginning of drug administration (Figure 6F,G).

**Figure 6.**
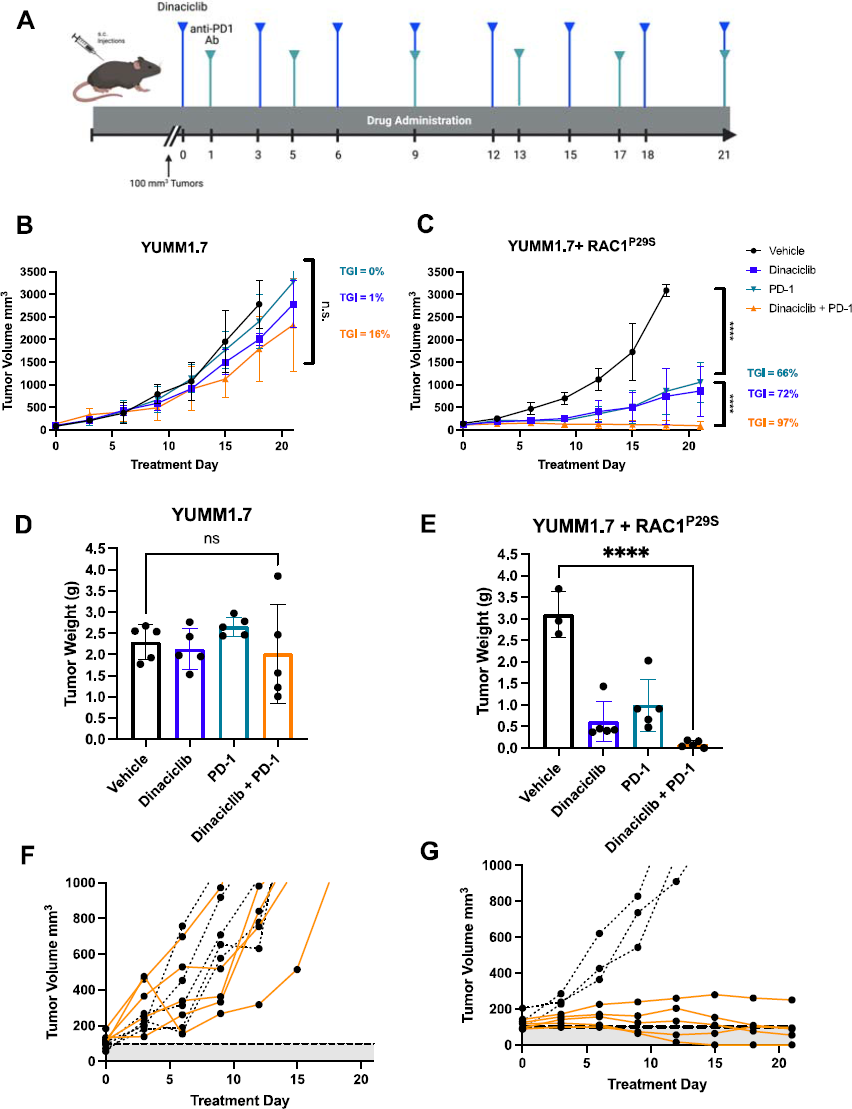
Combination therapy of CDK9 inhibition and anti-PD-1 immunotherapy significantly decreases tumor growth *in vivo*. **A.** Drug administration schedule. Mice were dosed with 30 mg/kg dinaciclib every 3 days and/or 10 mg/kg anti-PD1 Ab every 4 days via intraperitoneal injection (i.p.) **B.** Tumor volume by treatment day in YUMM1.7 and C. YUMM1.7 + RAC1^P29S^. Data points shown as mean with error. Tumor growth inhibition (TGI) calculated by (1-(mean treatment volume/mean vehicle volume))*100. **C.** Tumor volume of YUMM1.7 tumors and **E.** YUMM1.7 + RAC1 P^29S^ tumors. Bars represent the mean with standard deviation. **F-G.** Growth curves of each individual vehicle mouse (black) vs. dinaciclib and anti-PD1 (orange) combination. Dashed line at 100mm^3^ represents average tumor volume at day 0 of treatment.

### RAC1^P29S^ expression increases tumor immune cell infiltration

To determine the effect of the treatment regimen on tumor immune cell infiltration, we performed immunohistochemistry (IHC) on tumors fixed from the mice at the experiment endpoint. Overall, there was more T cell (CD8) and myeloid (CD11c) immune cell infiltration in RAC1^P29S^-expressing melanomas and expression of these was further increased upon treatment with dinaciclib (Figure 7A,B). Additionally, there was increased CD86 expression in the RAC1 mutant tumors, which is found on monocytes, activated B cells, and dendritic cells and is necessary for T cell activation [27] (Figure 7C). In contrast, in parental YUMM 1.7 grafts, neither dinaciclib nor anti-PD-1 Ab elicited an increase in CD8, CD11c, or CD86 (Figure 7A-C). Surprisingly, there was no significant difference in caspase-3 positivity in control vs. treated tumors, suggesting that either these cells cannot proliferate or that a non-apoptotic mode of programmed cell death is evoked by combined anti-CDK9 and anti-PD1 therapy.

**Figure 7.**
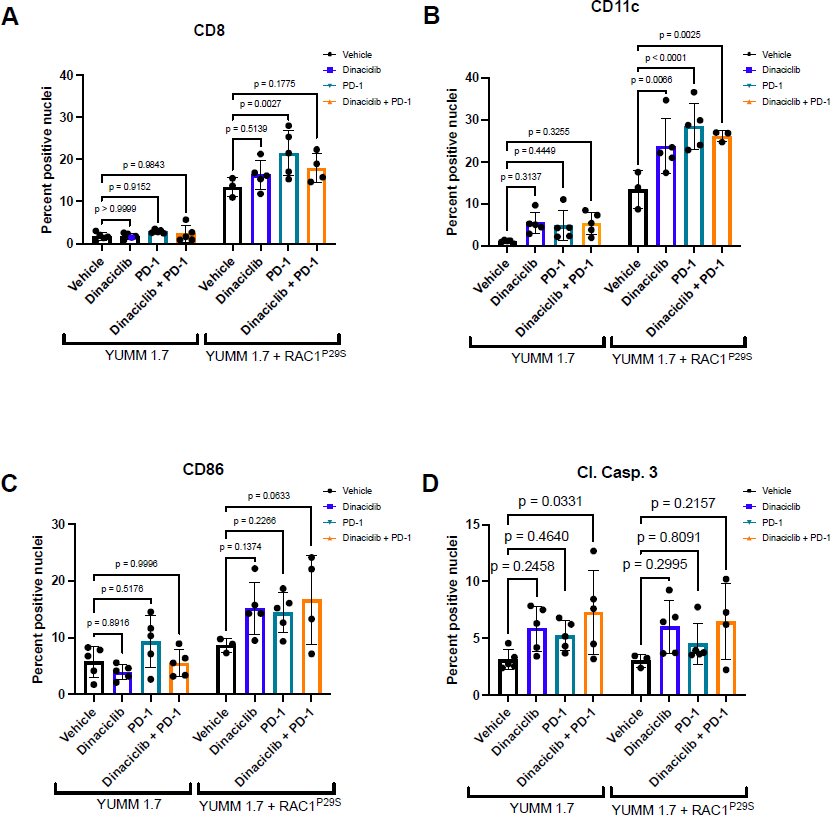
RAC1^P29S^ expression increases tumor immune cell infiltration. **A.** Percent positive nuclei from fixed tumor tissue at endpoint (day 21) of CD8, **B.** CD11c, **C.** CD86, and **D.** Cleaved caspase-3. P-values determined by multiple Student’s T-test relative to the vehicle.

## Discussion

*RAC1*-mutant melanomas display primary resistance to targeted BRAF and MEK inhibitors, higher metastatic propensity, and overall worse survival compared to *RAC1-* wild type melanomas [4, 5, 8]. There are currently no approved treatments specific for *RAC1*-mutant melanoma. Thus, there is a clinical need to identify novel targets that can combat this melanoma subtype or increase efficacy of existing treatments that would extend clinical benefits to a larger patient population. Immune checkpoint inhibitors have significantly improved patient prognosis in general in the field of melanoma, however, limited studies have investigated the RAC1 influence on sensitivity to these immune therapy agents. Previous studies have identified RAC1^P29S^-expressing melanocytes to express higher PD-L1 [9], indicating RAC1^P29S^ melanomas may be more sensitive to anti-PD-1 or PD-L1 ICI. Additional clinical data suggests systemic immunotherapies are beneficial to RAC1^P29S^ metastatic melanoma [10].

There have been recent mechanistic investigations into RAC1^P29S^-mediated primary BRAF and MEK inhibitor resistance. Mohan et al. demonstrate RAC1-driven elongation of lamellopodia sustains cell survival after treatment with MEK or BRAF inhibitors [28]. Furthermore, Lionarons et al. reported that RAC1^P29S^ drives an SRF/MRTF-mediated melanocytic to mesenchymal switch to promote BRAF inhibitor resistance. They further showed this resistance can be reversed with a SRF/MRTF inhibitor [29]. These works argue a role of actin cytoskeleton remodeling in propagation of BRAF inhibitor resistance. While our RNA-sequencing data was strongly enriched in cell cycle processes, which Lionarons et al. did not observe, our data uncovered many overlapping enriched gene sets, including TNFA_SIGNALING_VIA_NFKB, KRAS_SIGNALING_UP, and IL2_STAT5_SIGNALING [29].

The idea of employing CDK inhibitors to promote tumor immunogenicity has emerged over the past few years. These reports have established CDK inhibitors as pro-immunogenic by increasing both immune cell infiltration and increasing expression of molecules important in antigen presentation and T cell activation. Most of these works investigate combinations of CDK4/6 inhibitors with anti-PD-1 or anti-PD-L1 ICI [22–26]. In agreement with our data, additional prior reports demonstrated anti-tumor effects *in vivo* after combination of CDK9 inhibition via dinaciclib or other agents with anti PD-1 ICI [20, 21]. However, in *in vivo* models used by Zhang et al. and Hossain et al., *RAC1* wild type tumors benefited from CDK9 and anti-PD-1 combination, while only RAC1-mutant tumors were inhibited in our model. This could be explained by the high microsatellite instability status (MSI-H) of the engrafted tumor cells in the other studies. Supporting our observations, YUMM1.7 grafts have previously been shown to be insensitive to anti-PD-1 blockade [30]. Although PD-1 blockade combined with BRAF and MEK inhibition has demonstrated increased overall survival in *BRAF*-mutant melanoma, anti-PD-1 monotherapy response greatly varies in clinic [31]. Furthermore, contrary to Hossain et al., our findings indicate dinaciclib and PD-1 ICI-induced cell death is not solely immune-cell mediated in our syngeneic model [21]. This suggests an altered dependence of RAC1 mutant cells on CDK9, which could further be exploited for therapeutic intervention. The mechanism by which RAC1^P29S^ is linked to CDK9 remains unclear, however previous studies have alluded to a RAC1 influence on CDK9 activity, as RAC1 knockdown by siRNA resulted in decreased phosphorylation of RNA polymerase II at Ser 2 [32].

In this work, we establish CDK9 as a novel target in *RAC1*-mutant melanoma. Both non-transformed melanocytes and melanoma cells that overexpress RAC1^P29S^ and melanoma cell lines that harbor an endogenous *RAC1^P29S^* mutation displayed higher sensitivity to CDK9 inhibition by dinaciclib, NVP2, or a CRISPR-mediated CDK9 knockout. Inhibition of CDK9 induced higher surface expression of PD-L1 and MHC Class I *in vitro*. Combination of dinaciclib with anti-PD-1 Ab ICI resulted in tumor regression uniquely in syngeneic YUMM1.7 tumor grafts that express RAC1^P29S^. While few CDK9-specific inhibitors are currently in clinical trials, a potent multi-CDK inhibitor, such as dinaciclib, may offer additional anti-tumor benefits by inhibiting both the transcription regulating CDK9 and cell cycle regulating CDKs 1 and 2.

Collectively, these data indicate combination therapy of CDK9 inhibition with anti-PD-1 ICI may be particularly beneficial to melanoma patients that harbor the RAC1^P29S^ mutation. Furthermore, this work suggests the presence of RAC1-driven epigenetic reprograming events which might respond to epigenetic agents such as BRD4i. P-TEFb (CDK9-Cyclin T complex) binds to BRD4 and is subsequently recruited to promoter regions on chromosomes via BRD4’s interactions with acetylated histones [33, 34]. It has previously been reported that BRD4 binds to the promoter regions of cell cycle genes [35]. Furthermore, RNAi knockdown of BRD4 resulted in downregulation of genes involved in mitotic cell cycle, cell division, and DNA replication [36]. This could explain the strong cell cycle enrichment displayed in the RNA-seq and MIBs/MS. Since BRD4 is necessary for P-TEFb driven gene transcription, BRD4 may also yield a potential therapeutic target in *RAC1*-mutant melanoma. Several studies have shown promising anti-tumor effects by targeting BRD4 with JQ-1 in melanoma [37–40].

Relatedly, it has also been reported that JQ-1, along with other bromodomain and Extra-Terminal motif (BET) inhibitors, modulate the tumor immune microenvironment and sensitize the tumor to PD-1 or PD-L1 blockade [41, 42]. However, the efficacy of BRD4 inhibition in *RAC1*-mutant melanoma remains unexplored.

The benefits of combining CDK9 inhibition and anti-PD-1 immunotherapy in RAC-driven malignancies may extend beyond *RAC1^P29S^* melanoma. RAC1 is mutated in approximately 3% of head and neck squamous cell carcinomas (HNSCC), most commonly at the A159 (A159V) and P29 (P29S) loci [4, 5]. The current standard of care in HNSCC includes surgery and radiotherapy, although immunotherapies have been approved and are widely implemented [43]. In addition, HNSCCs typically bear a higher tumor mutational burden and immune infiltration relative to other cancer subtypes [44, 45]. These aspects make *RAC1-*mutant HNSCCs good candidates to benefit from immunotherapy, and combination with a CDK9 inhibitor may further enhance this efficacy. While the response of the *RAC1^A159V^* mutant to immunotherapy has yet to be investigated, it is presumed that A159V would behave similarly to the P29S mutation, as both are activating mutations that result in “fast-cycling” GTPases [46]. Moreover, both inhibition of RAC1 or depletion of CDK9 enhanced radiosensitivity in HNSCC cells in two independent studies, further implicating RAC1-mediated therapy resistance is influenced by CDK9 [47, 48].

Overall, these results demonstrate a novel treatment strategy for RAC1^P29S^-mutant melanoma. This therapeutic approach builds on the strategy of creating a “hotter” tumor microenvironment, which may improve clinical options for tumors that would not benefit from traditional treatment regimens.

## Authors’ Contributions

**A. Cannon:** Conceptualization, data curation, formal analysis, investigation, methodology, project administration, supervision, visualization, writing – original draft, writing – review and editing **K. Budagyan:** Conceptualization, formal analysis, investigation, writing – review and editing **C. Uribe-Alvarez:** Funding acquisition, methodology, writing – review and editing **A. Kurimchak:** Formal analysis, investigation, software **D. Araiza-Olivera:** Methodology **Q. Cai:** Formal analysis, investigation, methodology, software **S. Peri:** Data Curation, formal analysis, software **Y. Zhou:** Data Curation, formal analysis, software **J. Duncan:** Methodology, resources, supervision, formal analysis **J. Chernoff:** Conceptualization, funding acquisition, methodology, project administration, resources, supervision, visualization, writing – original draft, writing – review and editing

## Acknowledgments

We would like to thank Ruth Halaban and Gaudenz Danuser for cell lines. We would also like to thank Fox Chase Cancer Center core facilities including flow cytometry, biological imaging, laboratory animal and biostatistics facilities. This work was supported by grants from the NIH R01 CA2271844 (JC), the Melanoma Research Foundation (JC), the F.M. Kirby Foundation (CUA), NIH R01 CA211670 (JSD), NIH T32 CA009035 (AMK), and NCI Core Grant P30 CA06927 (to Fox Chase Cancer Center).

