## Supplemental Figures for "Unique vulnerability of *RAC1*-mutant melanoma to combined inhibition of CDK9 and immune checkpoints"

### Slide 1
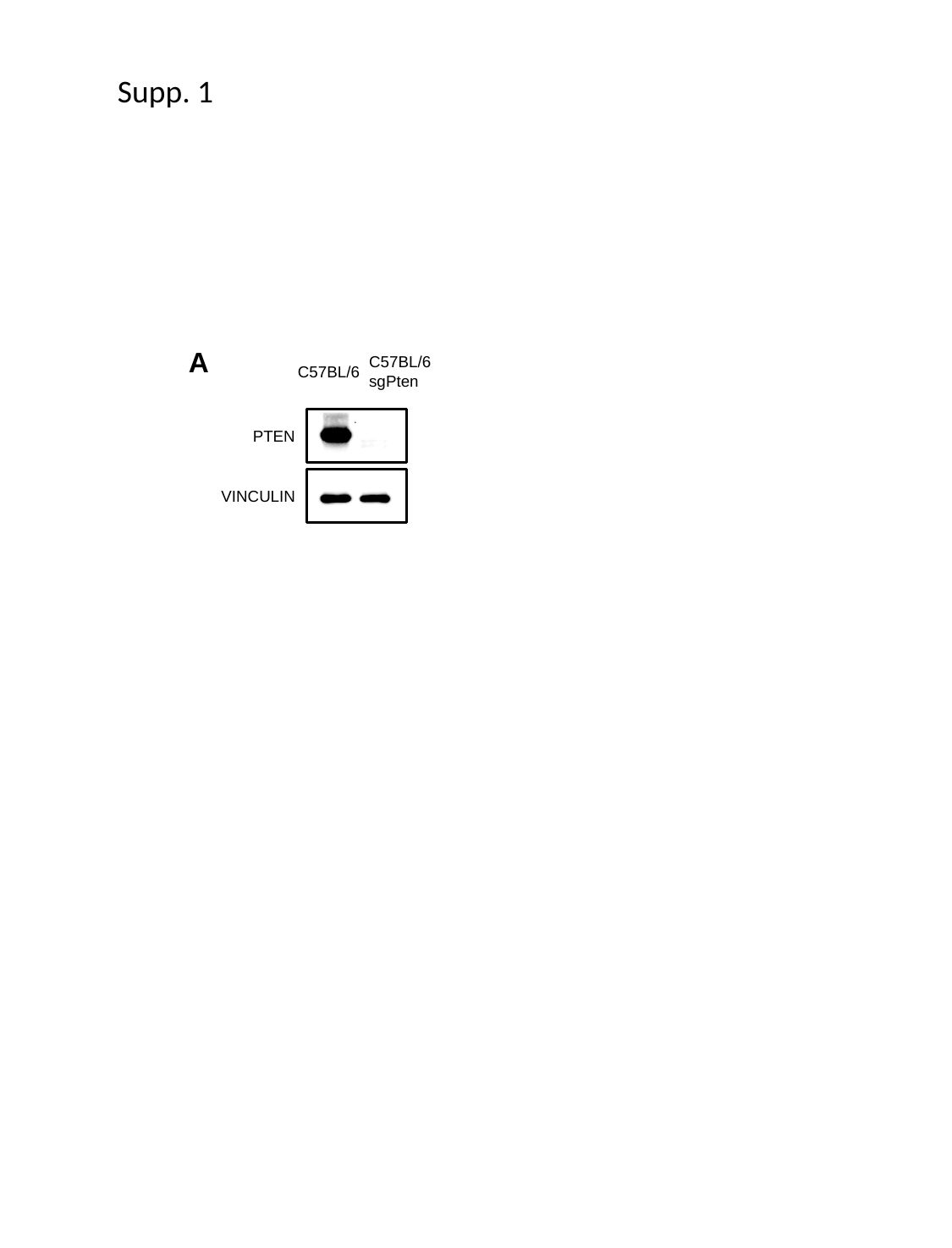

Supp. 1
A
C57BL/6 sgPten
C57BL/6
PTEN
VINCULIN

### Slide 2
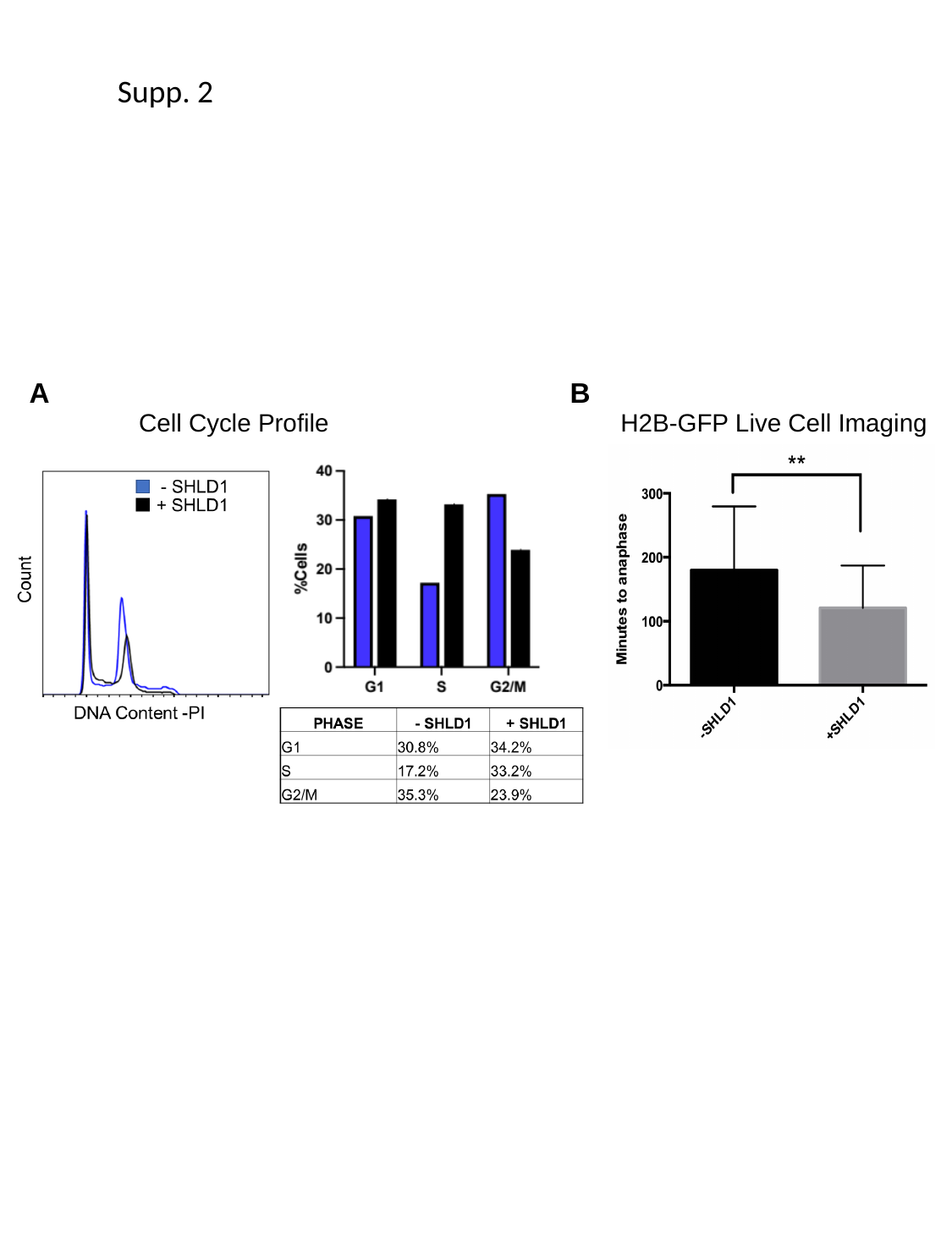

Supp. 2
A
B
Cell Cycle Profile
H2B-GFP Live Cell Imaging

### Slide 3
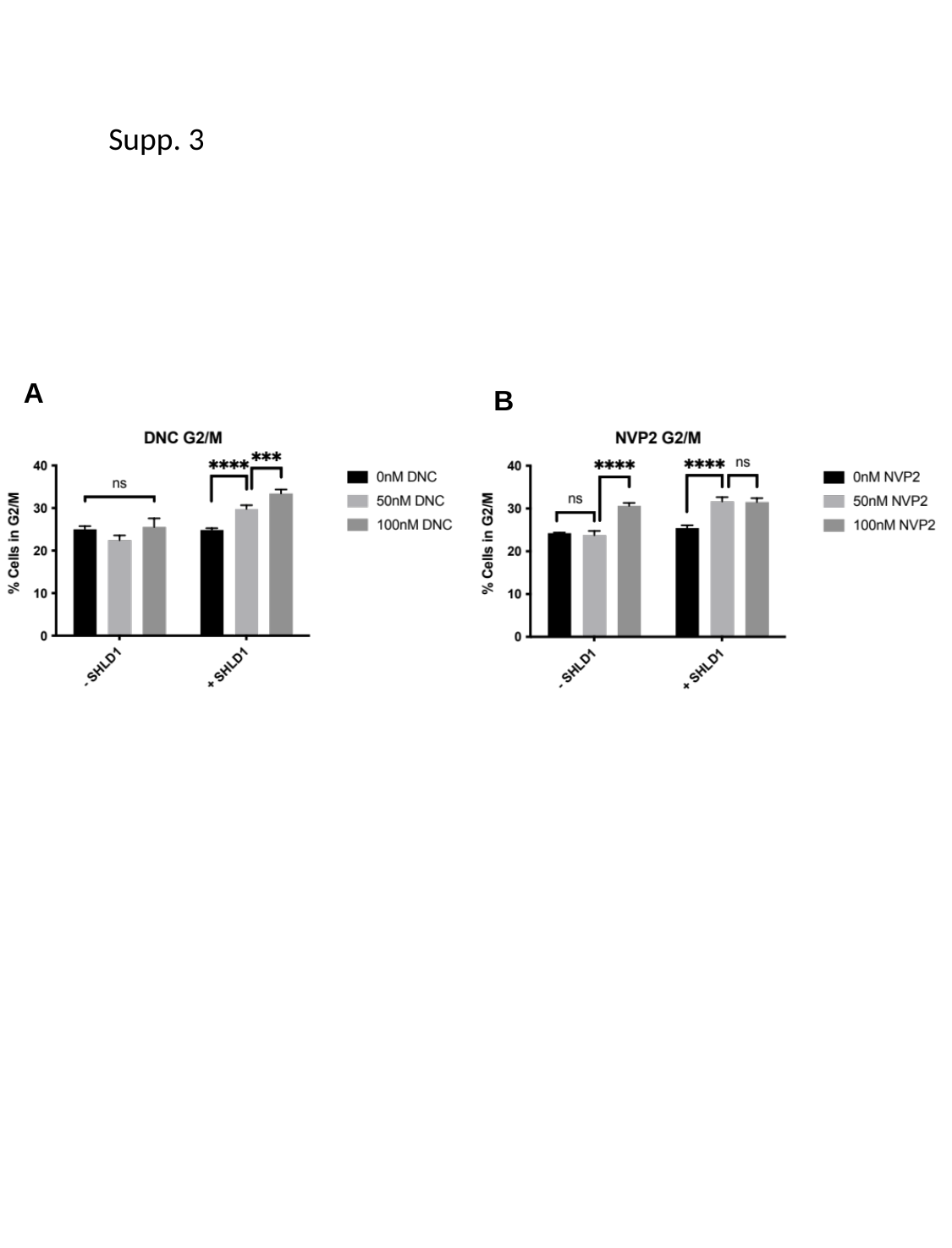

Supp. 3
A
B
